# Single-base m^6^A epitranscriptomics reveals novel HIV-1 host interaction targets in primary CD4^+^ T cells

**DOI:** 10.1101/2024.12.31.630958

**Authors:** Siyu Huang, Yutao Zhao, Stacia Phillips, Bethany Wilms, Chuan He, Li Wu

## Abstract

*N*^6^-methyladenosine (m^6^A) is the most prevalent cellular mRNA modification and plays a critical role in regulating RNA stability, localization, and gene expression. m^6^A modification plays a vital role in modulating the expression of viral and cellular genes during HIV-1 infection. HIV-1 infection increases cellular RNA m^6^A levels in many cell types, which facilitates HIV-1 replication and infectivity in target cells. However, the function of m^6^A modification in regulating HIV-1 infection of primary CD4^+^ T cells remains unclear. Here, we demonstrate that HIV-1 infection of Jurkat CD4^+^ T cells and primary CD4^+^ T cells promotes the interaction between the m^6^A writer complex subunits methyltransferase-like 3 and 14 (METTL3/METTL14). Using single-base m^6^A-specific RNA sequencing, we identified several differentially m^6^A-modified cellular mRNAs, including *perilipin 3* (*PLIN3*), during HIV-1 infection in primary CD4^+^ T cells. Interestingly, HIV-1 infection increased *PLIN3* mRNA level by enhancing its stability, but PLIN3 protein level was decreased. Knocking down PLIN3 in primary CD4^+^ T cells reduced HIV-1 production but enhanced virion infectivity. In contrast, in Jurkat cells, *PLIN3* mRNA and protein expression levels were unaffected by HIV-1 infection, and knocking out PLIN3 did not impact HIV-1 production or infectivity. These results indicate that the interplay between HIV-1 and PLIN3 is cell-type specific and only observed in primary CD4^+^ T cells. Overall, our results highlight the importance of m^6^A RNA modification in HIV-1-infected primary CD4^+^ T cells and suggest its significance as a regulatory mechanism in HIV-1 infection.

**Author Summary:** *N*^6^-methyladenosine (m^6^A) is a common chemical modification on mRNA that helps control RNA stability, localization, and gene expression. m^6^A modification of viral and cellular RNA is important for HIV-1 infection. In this study, we found that HIV-1 infection of CD4^+^ T cells enhanced the interaction between two proteins, METTL3 and METTL14, which are responsible for adding m^6^A modifications to RNA. Using m^6^A-specific RNA sequencing, we identified several mRNAs with altered m^6^A modifications during HIV-1 infection, including one called *PLIN3*. Interestingly, HIV-1 infection stabilized and increased *PLIN3* mRNA levels, but reduced PLIN3 protein expression in primary CD4^+^ T cells. When we knocked down PLIN3 in primary CD4^+^ T cells, it decreased HIV-1 production but made the HIV-1 particles more infectious. In contrast, in the Jurkat CD4^+^ T cell line, HIV-1 infection did not affect PLIN3 expression and knockout of PLIN3 did not alter HIV-1 production or infectivity, suggesting that the effect is specific to primary CD4^+^ T cells. Our findings show the importance of m^6^A RNA modification in HIV-1 infection by regulating host genes like *PLIN3* and suggest a unique regulatory mechanism in HIV-1 infected primary CD4^+^ T cells.

## Introduction

*N*^6^-methyladenosine (m^6^A) is the most prevalent modification found in eukaryotic RNA, and it reversibly regulates gene expression by influencing RNA stability, alternative splicing, and protein translation [1, 2]. This reversible modification is regulated by two groups of proteins involving a writer complex (methyltransferase) and erasers (demethylases). The m^6^A writer core complex consists of the catalytic subunit methyltransferase-like 3 (METTL3), and methyltransferase-like 14 (METTL14), which stabilizes METTL3 for substrate RNA binding [3–5]. m^6^A erasers includes fat mass and obesity-associated protein (FTO) and AlkB family member 5 (ALKBH5), which remove the methyl group [6, 7].

HIV-1 upregulates cellular m^6^A RNA levels in many target cell lines [8–12]. In addition, we reported that cellular RNA m^6^A level is increased in peripheral blood mononuclear cells (PBMCs) from HIV-1 infected patients, and this effect is reversed by antiretroviral therapy [13]. However, the mechanism of this cellular RNA m^6^A increase is not understood. Our group analyzed the expression levels of m^6^A writer and eraser proteins in primary CD4^+^ T cells from healthy donors [8] and HIV-1 latently infected J-Lat CD4^+^ T cells [12]. Based on these results, we hypothesize that the increase in cellular m^6^A levels is regulated at the level of writer complex formation or methyltransferase activity.

Several studies have identified m^6^A modifications on both cellular mRNA and HIV-1 RNA in various cell lines using m^6^A RNA Immunoprecipitation (meRIP) or crosslinking immunoprecipitation (CLIP) sequencing (reviewed in [14]). However, these methods are not of sufficient resolution to identify the exact site of m^6^A. Identifying the precise location of and quantitative changes in m^6^A modifications in specific cellular transcripts during HIV-1 infection is important for understanding how m^6^A modifications modulate cellular RNAs and HIV-1 infection. The m^6^A epitranscriptomic profile at single-base resolution has been reported for J-Lat CD4+ T cells under conditions of latency reversal [12]. However, it is important to define the m^6^A landscape in primary CD4^+^ T cells, which are the major target of HIV-1 infection *in vivo*.

In this study, we examined how HIV-1 infection affects cellular mRNA m^6^A levels and characterized the m^6^A epitranscriptomic profile at single-base resolution in HIV-1 infected primary CD4^+^ T cells. We show that HIV-1 increased cellular mRNA m^6^A levels and promoted the interaction between METTL3 and METTL14 in CD4^+^ T cells. Additionally, we identified specific changes in m^6^A modifications in a subset of cellular transcripts in HIV-1 infected primary CD4^+^ T cells compared to mock controls. The mRNA encoding for perilipin 3 (PLIN3) has significantly higher levels of m^6^A modification at a single site in HIV-1-infected cells compared to mock-infected controls. We go on to show that HIV-1 infection regulates *PLIN3* mRNA and protein levels, and knocking down PLIN3 reduces HIV-1 release but enhances virion infectivity in primary CD4^+^ T cells. These phenotypes are cell-specific and not observed in Jurkat T cells. These findings suggest that differential m^6^A modification is a key regulatory mechanism in HIV-1 infection and provide the basis for future functional studies into how differential m^6^A modification affects HIV-1 replication.

## Results

### HIV-1 upregulates m^6^A modification levels in cellular mRNA and promotes the interaction between METTL3 and METTL14 in CD4^+^ T cells

Our group has reported that HIV-1 infection upregulates m^6^A levels of total cellular RNA in CD4^+^ T cells without changing the expression of m^6^A writers and erasers [8, 12]. However, the mechanism of the m^6^A increase remains to be investigated. In the m^6^A writer complex, METTL3 is the catalytic subunit, while METTL14 binds to RNA substrate and stabilizes the m^6^A writer complex [4, 5]. Therefore, we hypothesized that the METTL3/14 interaction may be increased during HIV-1 infection. We first sought to determine the kinetics of m^6^A upregulation in response to HIV-1 infection of Jurkat cells to identify the optimal time of infection at which to measure the METTL3/14 interaction. As expected, the expression of HIV-1 proteins increased over a period of 120 hr (Fig 1A). Measurement of m^6^A levels in polyadenylated RNA showed that there was a similar increase in infected cells at both 72 and 120 hr post-infection (hpi) compared to mock-infected controls. (Fig 1B). Next, we tested whether the interaction between METTL3 and 14 would be changed by HIV-1 infection in Jurkat cells (Fig 1C). A METTL3 antibody was used for immunoprecipitation (IP) and IgG was used as a negative control. The results showed that there was a 3-fold increase in the amount of METTL14 that co-IPs with METTL3 in HIV-1-infected cells compared to mock-infected controls. We then sought to confirm these results in primary CD4^+^ T cells isolated from the PBMC of healthy blood donors. Viral infection was confirmed using p24 enzyme-linked immunosorbent assay (ELISA) (Fig 1D). m^6^A levels were measured and we observed a small but significant increase for each donor (Fig 1E). Total protein from these cells was used to perform co-IP as described above (Fig 1F). We again observed a ∼3-fold increase in the amount of METTL14 that co-Ips with METTL3 in HIV-1-infected cells compared to mock-infected controls. Overall, our results indicate that HIV-1 infection results in an increase in the interaction between METTL3 and 14 in CD4^+^ T cells that is associated with an increase in m^6^A levels.

**Fig 1.**
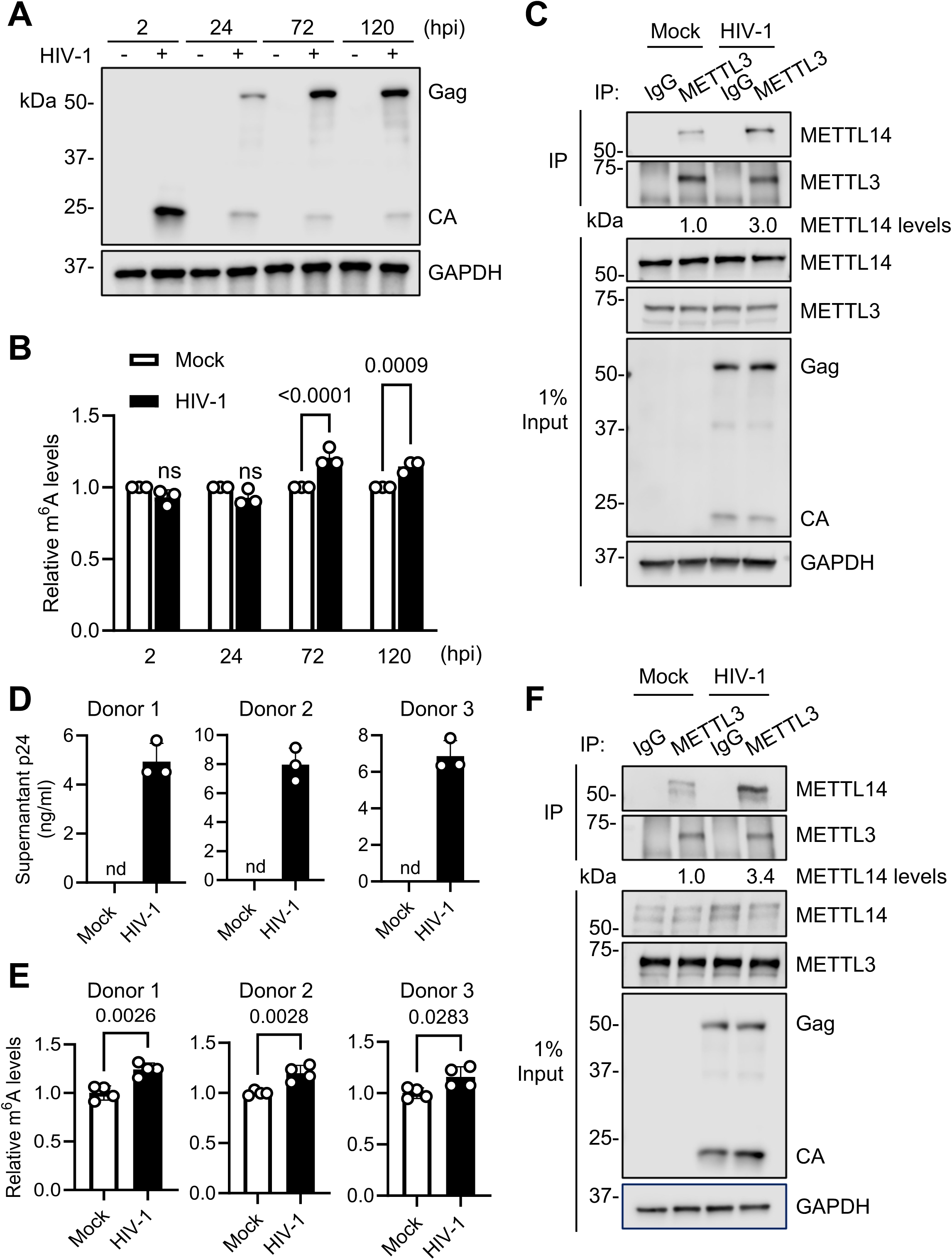
HIV-1 upregulates m^6^A modification levels in cellular mRNA and promotes the interaction between METTL3 and METTL14 in CD4^+^ T cells. **(A-C)** Jurkat cells were mock infected or infected with HIV-1_NL4–3_ at an MOI of 1. **(A)** Infection was confirmed by immunoblot (IB) analysis of HIV-1 Gag and capsid (CA) at the indicated times post-infection. **(B)** m^6^A levels in cellular mRNA from mock or HIV-1-infected cells were measured by ELISA at the indicated times post-infection. **(C)** Immunoprecipitation (IP) was performed at 72 hpi using non-specific IgG or an anti-METTL3 antibody. The indicated proteins were detected by IB in the input and IP samples. Relative levels of METTL14 in the IP were determined by densitometry (METTL14/METTL3). **(D-F)** Activated primary CD4^+^ T cells were mock-infected or infected with HIV-1_NL4–3_ at an MOI of 1 for 96 hr. **(D)** Infection was confirmed by measuring supernatant p24 levels by ELISA. nd, not detectable. **(E)** m^6^A levels in cellular mRNA from mock or HIV-1-infected cells were measured by ELISA. **(F)** IP was performed at using non-specific IgG or an anti-METTL3 antibody. The indicated proteins were detected by IB in the input and IP samples. Relative levels of METTL14 in the IP were determined by densitometry (METTL14/METTL3). Data are shown as mean ± SD from three individual experiments. Two-way ANOVA with Bonferroni correction (B) and two-tailed, unpaired *t*-test (E) were used for statistical analysis (*P* values are shown on figures). ns, not significant.

### m^6^A-SAC-Seq identifies m^6^A modifications in both cellular mRNAs and HIV-1 RNA

To identify the m^6^A sites in HIV-1 infected primary CD4^+^ T cells at single-base resolution, we used m^6^A-selective allyl chemical labeling and sequencing (m^6^A-SAC-seq) to identify m^6^A modifications at single-nucleotide resolution in primary cells [15, 16]. Primary CD4^+^ T cells were prepared from three individual healthy donors and infected with HIV-1 at a multiplicity of infection (MOI) of 1 for 96 hr. Polyadenylated RNA was purified with two rounds of poly(A)-enrichment prior to m^6^A-SAC-seq. Sequencing data was analyzed to identify individual m^6^A sites that are significantly changed in HIV-1-infected cells compared to mock controls. (Table S1). This analysis revealed 31,075 individual m^6^A modifications on 6,149 unique transcripts (Gene Expression Omnibus #280563). We created a heatmap to visualize and compare the m^6^A levels of cellular transcripts affected by HIV-1 infection (Fig 2A). Overall, 86 m^6^A sites became hypomethylated (blue) and 147 sites became hypermethylated (red) during HIV-1 infection (Fig 2B). Selected transcripts with a large log2 fold change (FC) or high degree of significance are indicated by arrows. Analysis of the transcript-level distribution of m^6^A sites shows that exons contain a major portion of m^6^A sites (46%), followed by 3’ UTR (38%), introns (14%), and 5’ UTR (2%) (Fig 2C). Analysis of the m^6^A motifs identified in cellular RNA reveal an internal consensus of GAC, with A/G/U and U/A/C in the terminal positions at the 5’ and 3’ ends of the motif, respectively (Fig 2D). To identify the pathways related to cellular transcripts that are hypermethylated during HIV-1 infection, we conducted gene ontology (GO) pathway analysis (Fig 2E). The analysis indicates that HIV-1 infection significantly regulates the m^6^A modification of transcripts involved in processes such as host RNA processing, signal transduction, and nucleocytoplasmic transport.

**Fig 2.**
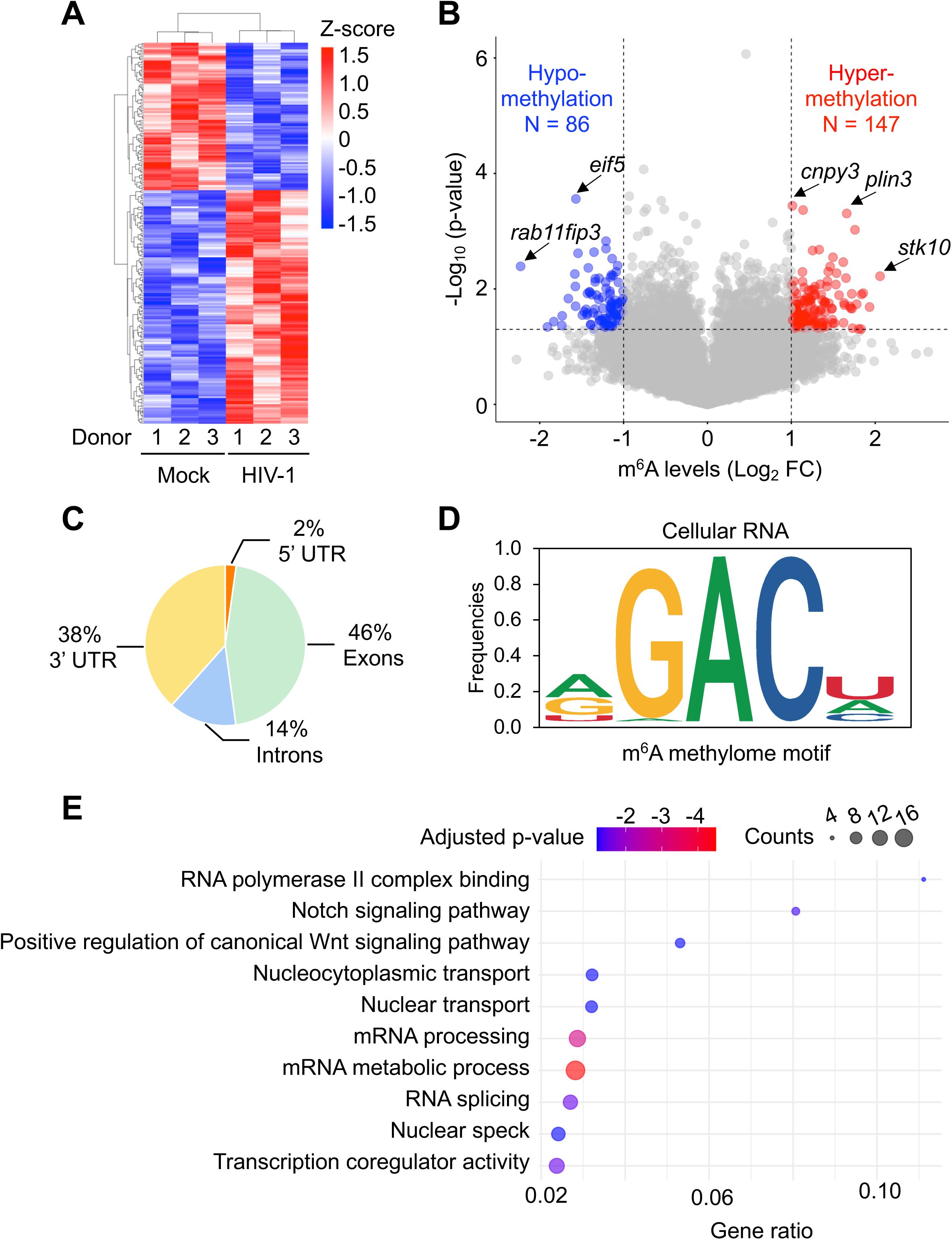
m^6^A-SAC-Seq identifies cellular mRNAs that are differentially m^6^A-modified upon HIV-1 infection. Activated primary CD4^+^ T cells isolated from donor PBMCs were mock-infected or infected with HIV-1_NL4–3_ at an MOI of 1 for 96 hr. Poly(A)-enriched RNA was used for m^6^A-SAC-seq. **(A)** Heat map showing transcript-level differences in m^6^A modification between mock and HIV-1-infected cells. Due to the large dataset, only genes with significant differences are displayed. Each row represents an RNA, and each column represents a sample. Both rows and columns are clustered using correlation distance. **(B)** Volcano plot showing m^6^A-hypomethylated (blue) and m^6^A-hypermethylated (red) mRNA from HIV-1 infected cells compared to mock-infected controls. Adenosines that are considered differentially methylated in response to HIV-1 infection are ≥ 2-fold changed compared to mock-infected controls, with *P* < 0.05. **(C)** m^6^A distribution in different regions of cellular mRNA. Analysis was performed with mock and HIV-1-infected samples combined (N = 6). **(D)** m^6^A consensus motif frequencies in cellular RNA were determined using m^6^A-SAC-seq. (**E)** Gene ontology (GO) analysis of m^6^A-hypermethylated cellular genes in Metascape. The top 10 pathways with the lowest adjusted p-values were selected and visualized using a bubble chart generated by R. Gene ratio is the percentage of genes in each GO term that are differentially changed. Adjusted p-value = Benjamini-Hochberg adjusted p-value.

We also identified 30 m^6^A sites in HIV-1 RNA from infected primary CD4^+^ T cells (Fig 3A. and Table S2). The m^6^A distribution in HIV-1 ORFs and the 3’UTR is shown in Fig 3B. Among these, the *pol* coding region was found to have the most m^6^A sites, with a total of 11. The site with the highest frequency of transcripts containing m^6^A modification is A8088, located in the sequence overlapping with the *env*, *rev, and tat* coding regions, with an average of 64.6% of transcripts having m^6^A modification among 3 individual donors. We further analyzed the m^6^A consensus motifs in viral RNA. We found that like cellular RNA, the internal consensus sequence in viral RNA was GAC (Fig 3C). However, m^6^A motifs in the viral RNA had a strong preference for A at the 5’ terminal position, in contrast to the more even distribution of A/G/U in cellular RNA m^6^A motifs. Moreover, there is a preference for U at the 3’ terminal position of m^6^A motifs in cellular RNA but not viral RNA. Overall, these data define the m^6^A epitranscriptomic landscape of cellular and viral RNA in HIV-1-infected primary CD4^+^ T cells.

**Fig 3.**
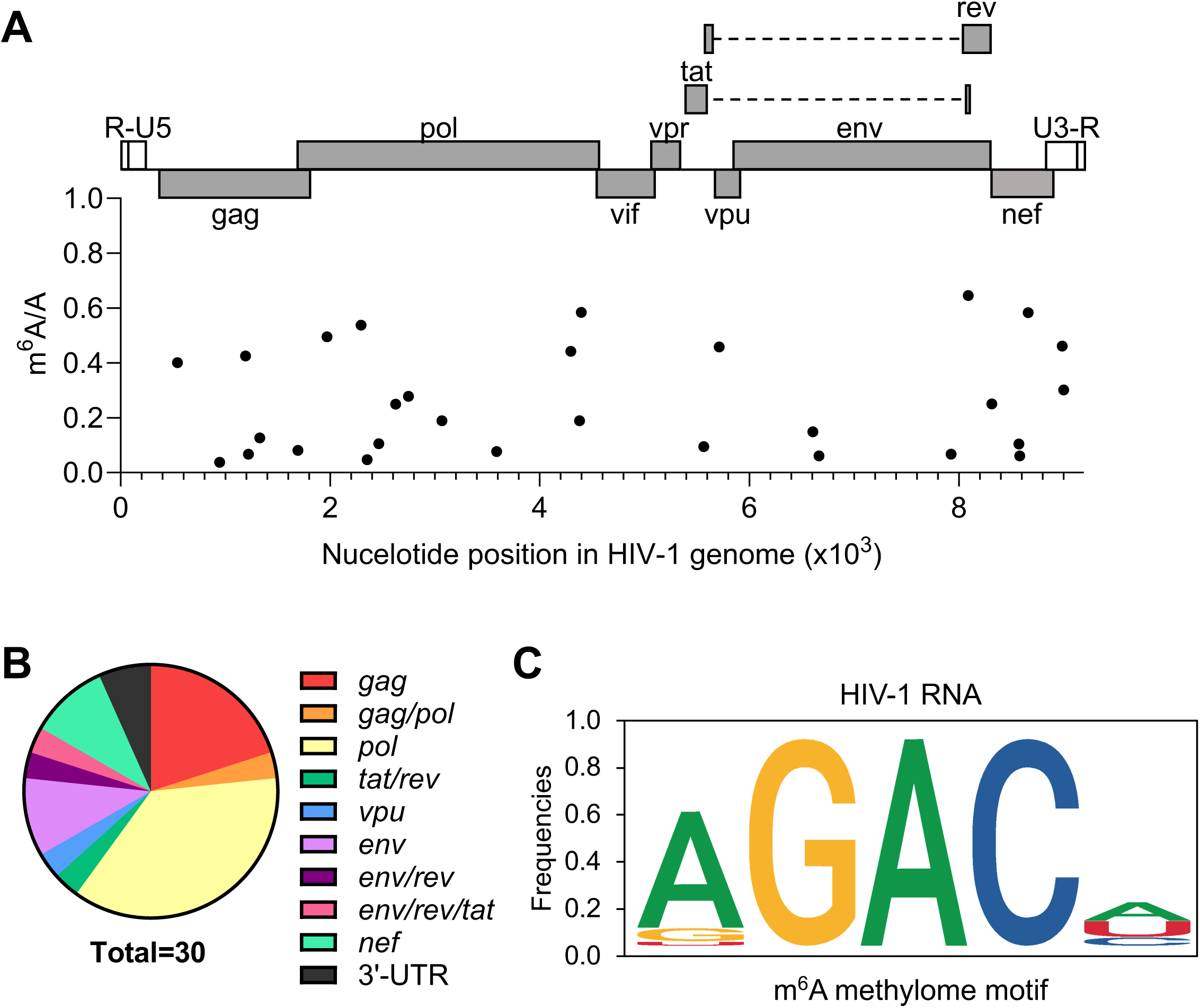
m^6^A distribution in HIV-1 genomic RNA. **(A)** HIV-1 RNA m^6^A sites and their frequencies are mapped to their nucleotide position in the HIV-1 genome (GenBank: AF033819.3) **(B)** Distribution of m^6^A sites in HIV-1 RNA. **(C)** m^6^A consensus motif frequencies in HIV-1 RNA were determined using m^6^A-SAC-seq.

### m^6^A modification of *PLIN3* mRNA is increased by HIV-1 infection in CD4^+^ T cells

Because most m^6^A modifications in our data set were located in exons, we selected 10 genes with significantly hypermethylated m^6^A sites located in an exon to validate our m^6^A-SAC-seq results (Table 1). We then performed meRIP to measure the relative m^6^A levels on the selected transcripts in HIV-1 infected Jurkat cells compared to mock-infected controls (Fig 4A). The results show that among these 10 genes, only the m^6^A levels of disco interacting protein 2 homolog B (*DIP2B*), *PLIN3*, and MYC binding protein 2 (*MYCBP2*) showed a global increase in m^6^A levels in HIV-1-infected cells. The observed hypermethylation of *PLIN3* mRNA is highly reproducible among primary cell donors (Table 1). Therefore, we chose *PLIN3* mRNA for further validation and functional studies. meRIP was used to confirm a significant increase in *PLIN3* mRNA m^6^A levels in HIV-1-infected primary CD4^+^ T cells compared to mock-infected controls (Fig 4B, C).

**Fig 4.**
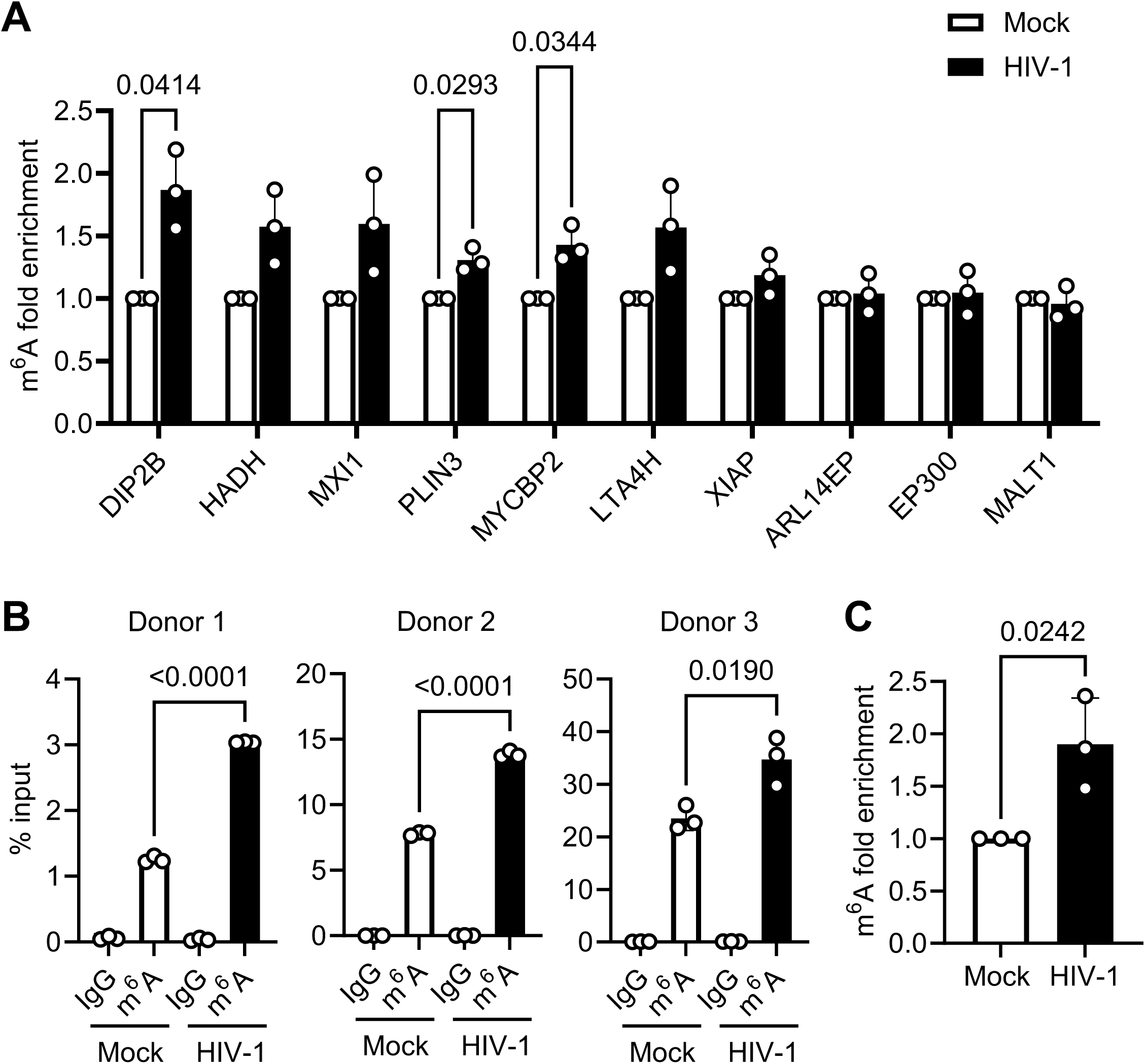
m^6^A modification of PLIN3 mRNA is increased by HIV-1 infection in CD4^+^ T cells. **(A)** Jurkat cells were mock-infected or infected with HIV-1_NL4–3_ at an MOI of 1 for 72 hr. Total cellular RNA was subjected to meRIP, and the enrichment of m^6^A-modified transcripts in the m^6^A-IP was determined relative to mock-infected controls. **(B-C)** Activated primary CD4^+^ T cells isolated from donor PBMCs were mock-infected or infected with HIV-1_NL4–3_ at an MOI of 1 for 96 hr. Total cellular RNA was subjected to meRIP, and the level m^6^A-modified transcripts in the m^6^A-IP was determined relative to **(B)** input or **(C)** mock-infected controls. Data are shown as mean ± SD. Multiple unpaired *t*-test (A) or two-tailed unpaired *t*-test (B, C) were used for statistical analysis (*P* values are shown on figures).

**Table 1.**
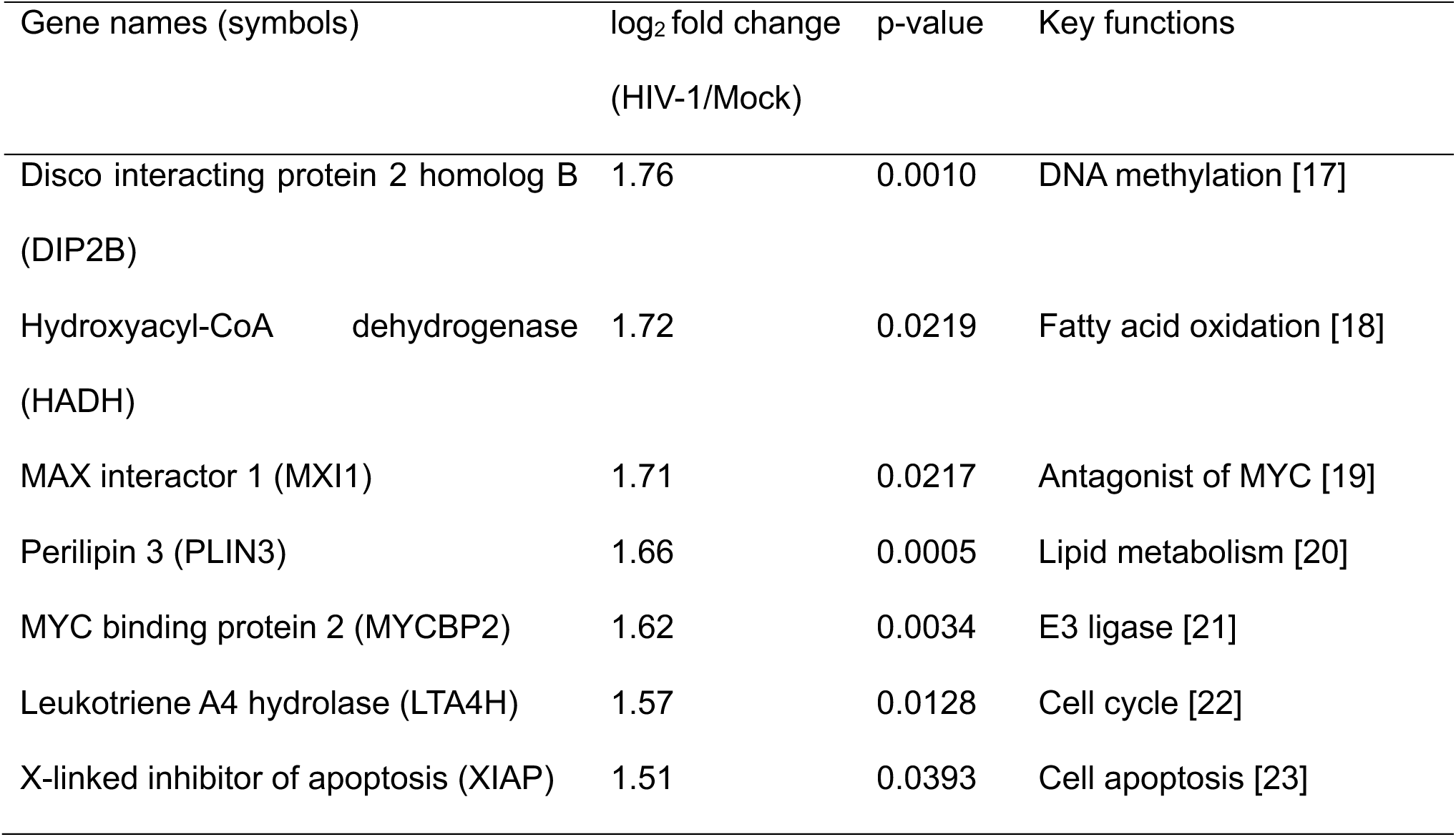
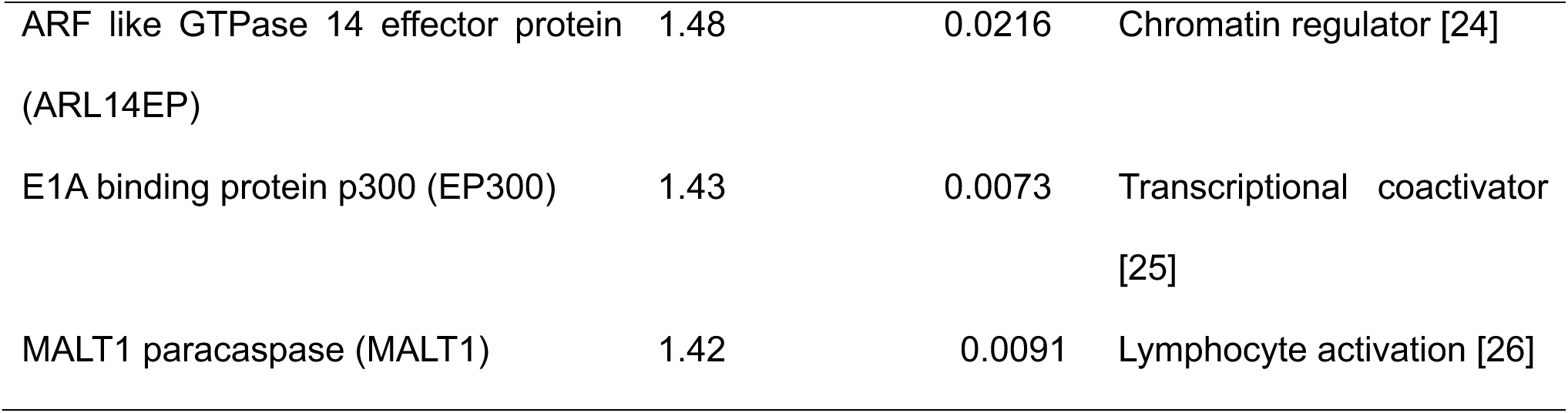
Top 10 cellular transcripts with significant m6A hypermethylation in HIV-1 infected primary CD4+ T cells. **note.** Activated primary CD4+ cells from three different healthy donors were mock-infected or infected with HIV-1NL4-3 at an MOI of 1 for 96 hr. Poly(A)-enriched RNA from cells was analyzed by m6A-SAC-seq. The top 10 cellular genes with significant m6A hypermethylation are listed in order of log2 fold change in m6A levels (p < 0.05). All cellular transcripts analyzed by m6A-SAC-seq are included in Table S1 (Excel file).

### PLIN3 does not affect HIV-1 replication in Jurkat cells

Based on our findings that HIV-1 infection promotes m^6^A methylation of *PLIN3* mRNA in CD4^+^ T cells, we asked whether infection also changes PLIN3 expression in Jurkat CD4^+^ T cells. First, we used HIV-1 to infect Jurkat cells at an MOI of 1 for 72 hr and then detected *PLIN3* mRNA and protein levels by qRT-PCR and immunoblotting, respectively. We found that HIV-1 infection did not change the steady state level of *PLIN3* mRNA in Jurkat cells (Fig 5A). Likewise, PLIN3 protein levels also remain unchanged after HIV-1 infection of Jurkat cells (Figs. 5B and C). We next aimed to determine whether PLIN3 regulates HIV-1 replication in Jurkat cells. To test this, we generated a *PLIN3* knockout (KO) Jurkat cell line by lentiviral transduction of vectors expressing Cas9 and PLIN3 sgRNA. Jurkat cells transduced with sgScramble were used as control (Ctrl). Ctrl and PLIN3 KO cells were infected with HIV-1 at an MOI of 1 for 72 hr. Immunoblotting was used to determine the expression levels of PLIN3 and HIV-1 proteins (Fig 5D). As expected, no PLIN3 expression was observed in *PLIN3* KO cells and there was no difference in PLIN3 expression in Ctrl cells after HIV-1 infection. Further, we observed no difference in the expression of HIV-1 Env, Gag, or CA (Fig 5D, E). We measured HIV-1 p24 release in cell culture supernatants and found no difference between the Ctrl and PLIN3 KO cells, suggesting that PLIN3 is not required for virion release. Finally, virus input was normalized by p24 content prior to infection of TZM-bl cells to measure viral infectivity. The results confirm that the infectivity of HIV-1 virions produced in *PLIN3* KO cells was not changed from that in Ctrl cells (Fig 5G). Overall, our results demonstrate that HIV-1 infection does not alter the RNA or protein levels of PLIN3, and PLIN3 KO does not change HIV-1 infection in Jurkat CD4^+^ T cells. These results are consistent with a previous report of PLIN3 and HIV-1 infection [27].

**Fig 5.**
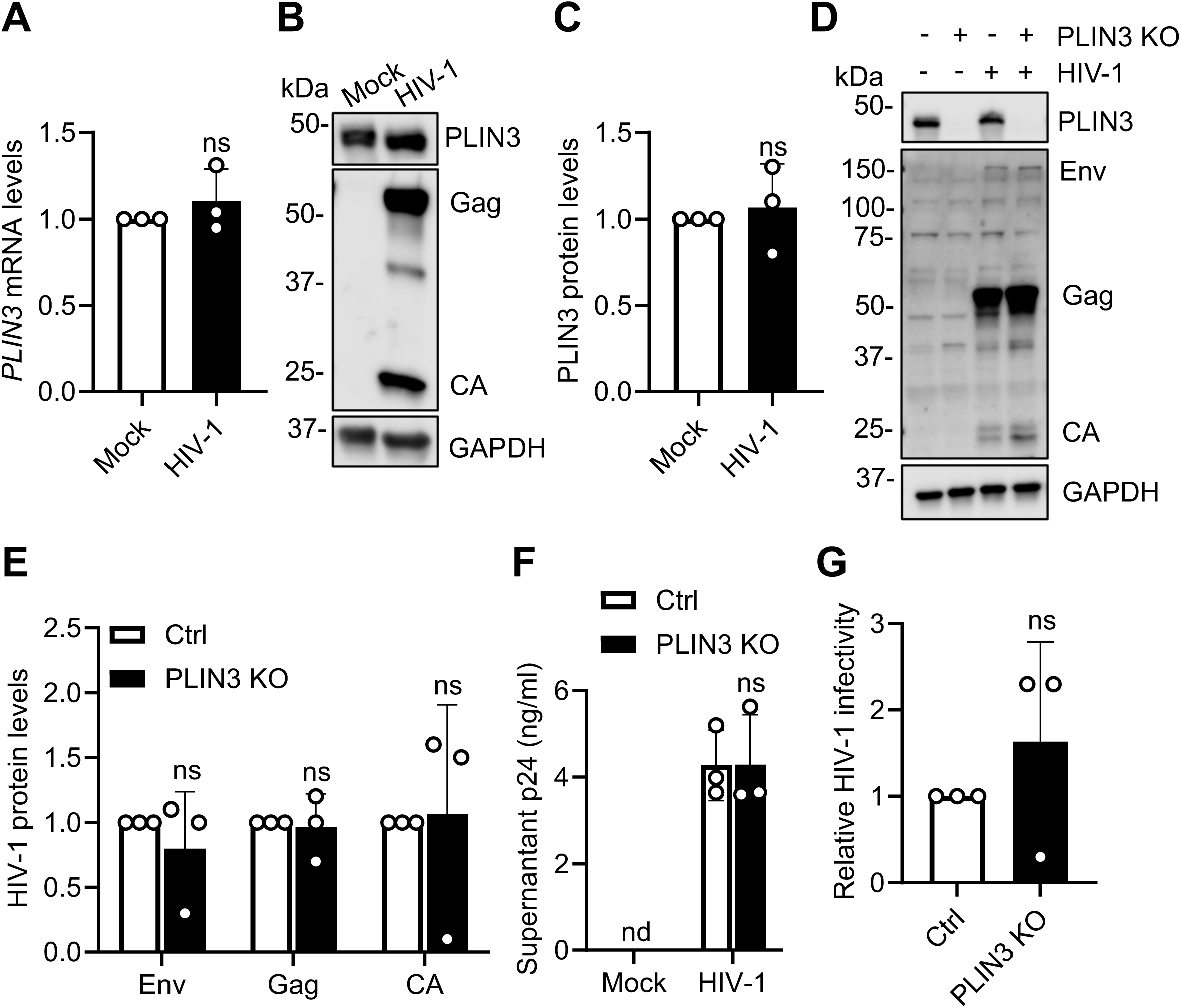
PLIN3 does not affect HIV-1 replication in Jurkat cells. **(A-C)** Jurkat cells were mock-infected or infected with HIV-1_NL4–3_ at an MOI of 1 for 72 hr. **(A)** PLIN3 mRNA levels were measured by qRT-PCR. **(B)** PLIN3 and HIV-1 protein expression was measured by IB. A representative IB is shown. **(C)** Relative quantification of PLIN3 protein expression as shown in (B) from three individual experiments. **(D)** Control (Ctrl) and PLIN3KO Jurkat cells were mock-infected or infected with HIV-1_NL4–3_ at an MOI of 1 for 72 hr. PLIN3 and HIV-1 protein expression was measured by IB. One individual experiment result is shown. **(E)** Relative quantification of HIV-1 protein expression as shown in (D) from three individual experiments. **(F)** Cell culture supernatants were collected from Ctrl and PLIN3 KO Jurkat cells with and without HIV-1 infection, and p24 levels were quantified by ELISA. nd, not detectable. **(G)** TZM-bl cells were infected with HIV-1 collected from Ctrl or PLIN3 KO cell culture supernatant. Luciferase activity was measured at 48 hpi. Data are shown as mean ± SD from three individual experiments. Two-tailed, unpaired *t*-test (A, C, F, and G) and multiple unpaired *t*-test (E) were used for statistical analysis. ns, not significant.

### HIV-1 infection increases *PLIN3* mRNA levels by enhancing *PLIN3* mRNA stability in primary CD4^+^ T cells

We next sought to determine whether HIV-1 infection alters the RNA or protein levels of *PLIN3* in primary CD4^+^ T cells. Primary CD4^+^ T cells were mock-infected or infected with HIV-1 at an MOI of 1 for 96 hr. To block HIV-1 replication, we used the reverse transcription inhibitor nevirapine (NVP) as a control. HIV-1 Gag and CA were detected to confirm infection. Immunoblotting showed that the levels of PLIN3 was significantly decreased by HIV-1 infection (Fig 6A, B). In contrast, the levels of *PLIN3* mRNA were increased by HIV-1 infection (Fig 6C).

**Fig 6.**
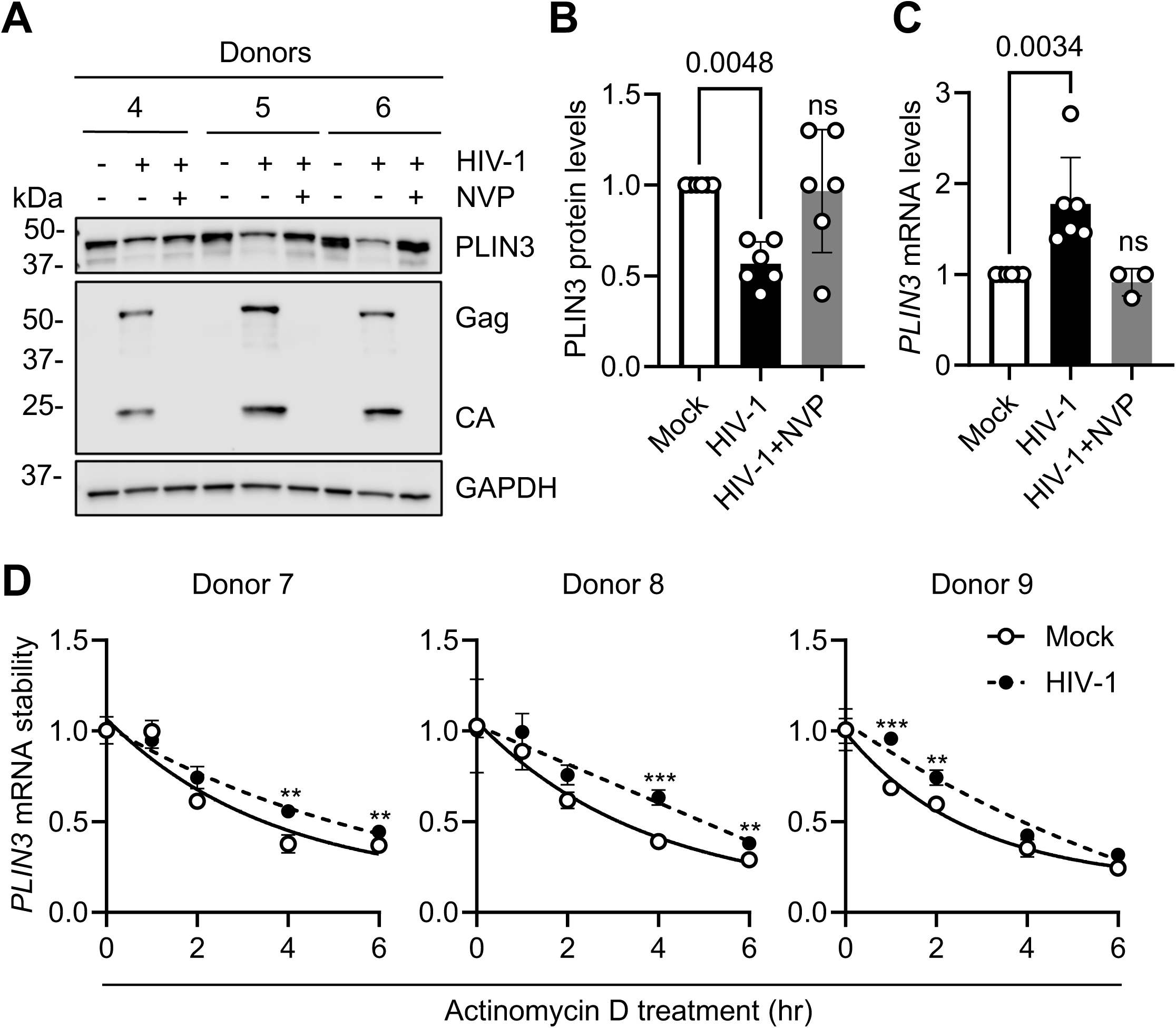
HIV-1 infection increases PLIN3 mRNA levels by enhancing PLIN3 mRNA stability in primary CD4+ T cells. **(A-C)** Primary CD4^+^ T cells were mock-infected or infected with HIV-1_NL4– 3_ at an MOI of 1 for 96 hr. The HIV-1 reverse transcription inhibitor NVP was used to block viral replication (HIV-1+NVP). **(A)** PLIN3 and HIV-1 protein expression was measured by IB. A representative IB is shown. **(B)** Relative quantification of PLIN3 protein expression as shown in (A) from three individual donors. **(C)** PLIN3 mRNA levels were measured by qRT-PCR. N = 6 (Mock, HIV-1) or N = 3 (HIV-1 + NVP). **(D)** Cells were treated with actinomycin D at 96 hpi. Samples were collected at the indicated time points, and PLIN3 mRNA levels were detected by qRT-PCR. Data are shown as means ± SD. Ordinary One-way ANOVA with Dunnett correction (B, and C) and multiple unpaired *t*-test (D) were used for statistical analysis (*P* values are shown on figures). ns, not significant. ** *P* < 0.01. *** *P* < 0.001.

Given that m^6^A methylation typically regulates mRNA levels by reducing mRNA stability [28, 29], we examined *PLIN3* mRNA stability in mock and HIV-1 infected primary CD4^+^ T cells. Cells were mock-infected or infected with HIV-1 at an MOI of 1 for 96 hr. Actinomycin D was then added to the culture medium to inhibit transcription, and samples were collected at the indicated times to measure the relative levels of *PLIN3* mRNA. The results indicate that *PLIN3* mRNA is more stable in HIV-1 infected cells compared to mock (Fig 6D), which may explain why the level of *PLIN3* mRNA level is higher in HIV-1 infected cells.

### Knockdown of PLIN3 in primary CD4^+^ T cells decreases HIV-1 production but increases viral infectivity

We next sought to investigate whether PLIN3 is necessary for HIV-1 infection in primary CD4^+^ T cells. Primary CD4^+^ T cells were transduced with lentiviral vectors expressing Cas9 and Scramble (sgCtrl) or PLIN3 (sgPLIN3), followed by infection with HIV-1 at an MOI of 1 for 96 hr. Immunoblot analysis indicated that PLIN3 levels were reduced by ∼2-fold in cells from all three donors (Fig 7A, B). PLIN3 did not affect the levels of cell-associated Env, Gag, or CA (Fig 7A, C). p24 levels in the cell culture supernatants were quantified and found to be reduced in sgPLIN3 cells compared to sgCtrl (Fig 7D). Virus input was normalized by p24 content prior to infection of TZM-bl cells to measure viral infectivity. The results showed that the infectivity of HIV-1 produced by sgPLIN3 cells is significantly increased compared to sgCtrl cells (Fig 7E). These results suggest that PLIN3 positively regulates HIV-1 production but inversely affects viral infectivity.

**Fig 7.**
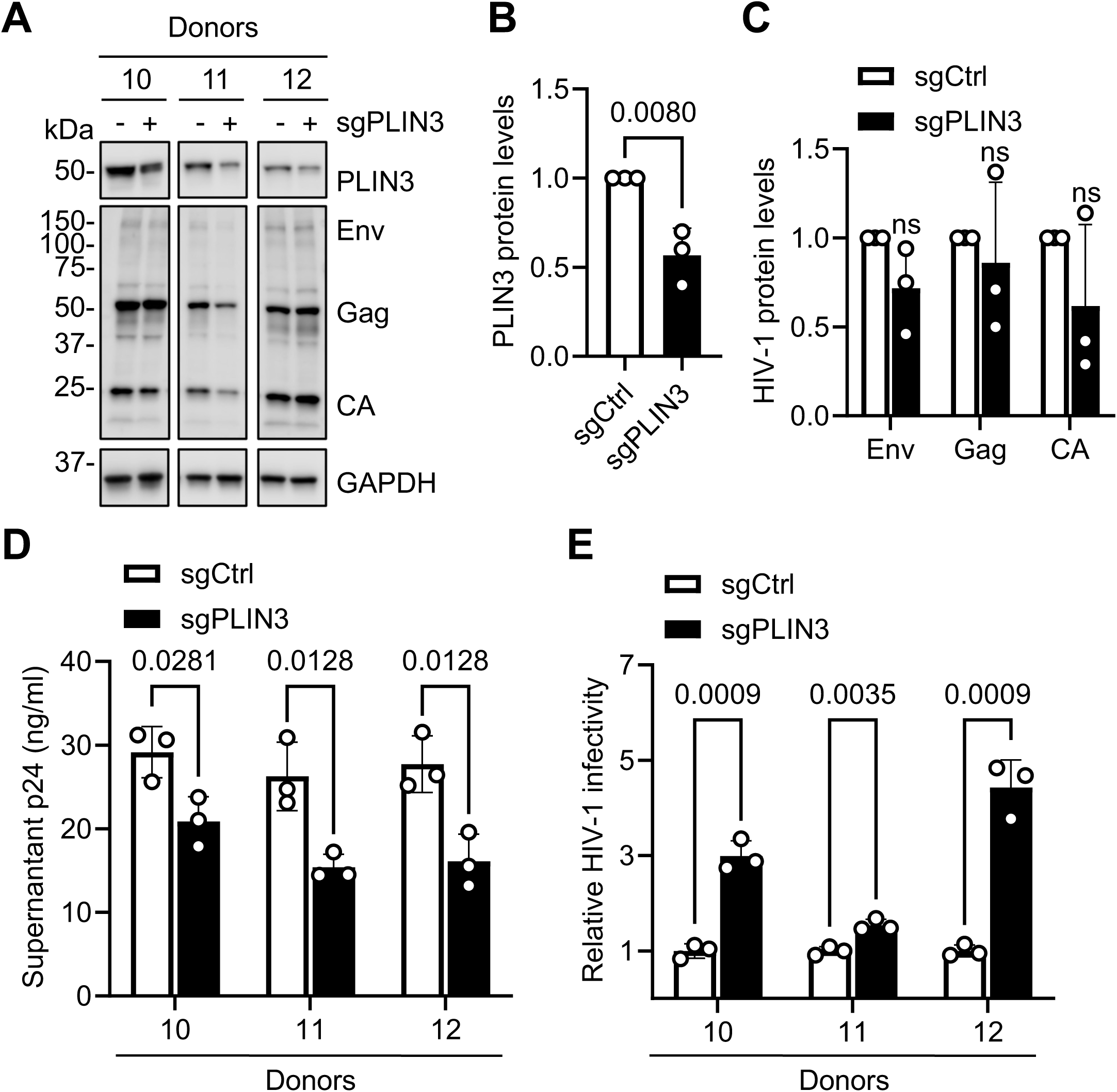
Knockdown of PLIN3 in primary CD4^+^ T cells decreases HIV-1 production but increases viral infectivity. **(A-D)** Primary CD4^+^ T cells were transduced with lentiviral vectors expressing non-targeting (Ctrl) or PLIN3 small guide (sg) RNA to achieve partial stable knockdown of PLIN3. Cells were then infected with HIV-1_NL4–3_ at an MOI of 1 for 96 hr. **(A)** Relative levels of PLIN3 expression and HIV-1 infection were measured by IB in cells from three independent donors. **(B)** Relative quantification of PLIN3 protein expression shown in (A). **(C)** Relative levels of HIV-1 protein expression shown in (A). **(D)** Cell supernatant p24 levels from HIV-1 infected cells were quantified by ELISA. **(E)** TZM-bl cells were infected with HIV-1 collected from sgCtrl or sgPLIN3 cell culture supernatants. Luciferase activity was measured at 48 hpi. Data are shown as means ± SD from three individual donors. Two-tailed unpaired *t*-test (B) and multiple unpaired *t*-test (C, D, and E) were used for statistical analysis (*P* values are shown on figures). ns, not significant.

In summary, this study provides new insight into the m^6^A epitranscriptomic landscape in HIV-1 infected primary CD4^+^ T cells and highlights a novel role for PLIN3 as a regulator of HIV-1 infection in a cell-type specific manner (Fig 8).

**Fig 8.**
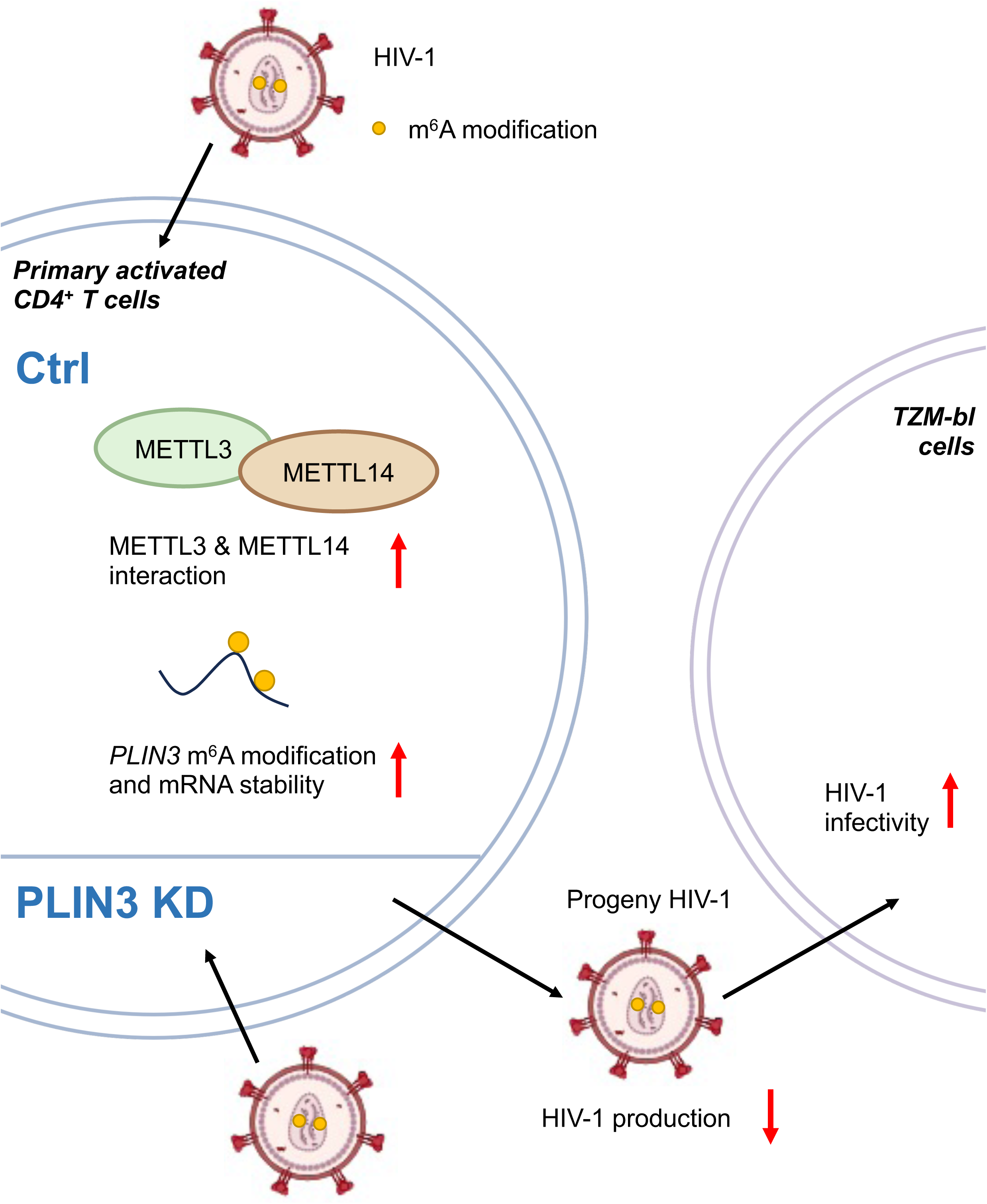
Summary and proposed model. In primary CD4^+^ T cells, HIV-1 infection promotes the interaction between METTL3/METTL14. HIV-1 infection increases PLIN3 m^6^A level and RNA stability. Knockdown of PLIN3 in primary CD4^+^ T cells decreases HIV-1 production but increases viral infectivity in TZM-bl cells. Ctrl, control; KD, knockdown.

## Discussion

m^6^A modification of HIV-1 RNA and the proteins involved in deposition, recognition, and removal of m^6^A marks play important roles in HIV-1 replication by regulating RNA stability, alternative splicing, RNA packaging, and Gag synthesis [30–34]. In this study, we sought to identifying cellular transcripts that are differentially m^6^A modified upon HIV-1 infection to provide novel insight into how HIV-1 regulates host gene expression.

It is well-established that HIV-1 infection causes an upregulation of cellular RNA m^6^A levels in a variety of cell types [8–12]. These findings suggest that HIV-1 may exploit the host m^6^A machinery to modulate viral infection. Alternatively, the increase in m^6^A may be a general host cell response to HIV-1 infection. Regardless, how m^6^A upregulation occurs during HIV-1 infection remains unclear. Our previous results showed that the expression levels of m^6^A writers and erasers were not altered by HIV-1 infection in primary CD4^+^ T cells or latently infected cells after reactivation [8, 12]. The absence of changes in writer or eraser protein levels suggests that the upregulation of m^6^A may result from an increase in methyltransferase activity rather than protein expression. Therefore, we conducted co-IP to examine the level of interaction between METTL3 and 14 during HIV-1 infection compared to uninfected controls. We found that HIV-1 infection enhances the METTL3/14 interaction in CD4^+^ T cells (Fig 1C, F). The increased interaction could potentially be attributed to post-translational modifications (PTMs) of METTL3 or METTL14. Another possible mechanism of m^6^A upregulation is PTM of m^6^A writers or erasers which can regulate protein stability or enzyme activity, thereby influencing the overall dynamics of m^6^A regulation [35, 36].

Previously reported m^6^A sequencing of RNA from HIV-1-infected cells using meRIP-seq and CLIP-seq has provided valuable insights into the location of RNA m^6^A modifications [37–39]. However, two major disadvantages of these methods are low resolution and the lack of m^6^A/A quantification. In this study, we employed m^6^A-SAC-seq to quantitatively identify individual m^6^A modifications on a transcriptome-wide scale in both cellular and HIV-1 RNA [15, 16]. These data are the first to report how productive HIV-1 infection regulates m^6^A modification, at single-base resolution, in primary CD4^+^ T cells and provide a foundation for targeted functional studies.

For the purposes of the current study, we chose to focus on transcripts that become significantly hypermethylated in primary CD4^+^ T cells upon HIV-1 infected compared to mock-infected controls. Our GO pathway analysis of these transcripts found an association between HIV-1 infection and mRNA splicing (Fig 2E), which is consistent with a previous study that performed gene set enrichment analysis (GSEA) of m^6^A sequencing from HIV-1 infected hippocampus from a transgenic rat [40]. This suggests that m^6^A modification of host cell RNA may be a regulatory mechanism of gene expression that affects RNA splicing during HIV-1 infection.

We identified a total 30 m^6^A modifications in HIV RNA, which is fewer than our previous analysis of RNA from J-Lat cells grown under conditions of latency reactivation. It is possible that HIV-1 RNA expressed after latency reactivation is more heavily m^6^A-modified than transcripts made during productive infection [12]. However, given the overlap between these two data sets (22 out of 30 m^6^A sites) it is more likely that this difference reflects the much lower percentage of primary cells expressing HIV-1 transcripts (data not shown). Of the 8 m^6^A sites unique to the current study, 7 were present on less than 10% of transcripts (Table S2). However, site A8660 in the *nef* region was modified in 58.3% transcripts, making this an interesting m^6^A modification for further study. Consistent with our previous study, three high frequency modification were present at A8088, A8984, and A8998 (Fig 3A, and Table S2). These three m^6^A sites were also identified by DRS in both HIV-1 producer HEK293T cells and infected CD4^+^ T cells and were implicated in viral RNA splicing [12, 31].

We chose to focus further on m^6^A modification of *PLIN3* mRNA during HIV-1 infection of primary CD4^+^ T cells. PLIN3, also known as TIP47 (Tail-interacting protein of 47 kDa), is a protein that plays a crucial role in lipid droplet formation [20]. It has been demonstrated that lipid rafts are important for the replication of many viruses in multiple cell types [41]. Particularly, plasma membrane rafts and HIV-1 Gag interaction play a critical role in HIV-1 assembly and release HIV-1 [42]. While there is no publication has reported that PLIN3’s expression level is altered by HIV-1 infection in primary CD4^+^ T cells, this study is the first one to reveal the effects of HIV-1 infection on PLIN3 RNA methylation and expression levels in primary CD4^+^ T cells. Interestingly, in primary cells, HIV-1 infection not only increased PLIN3 m^6^A methylation but also promoted its mRNA stability, resulting in an upregulation of *PLIN3* mRNA level. A previous study identified that insulin-like growth factor 2 mRNA-binding proteins (IGF2BPs; including IGF2BP1/2/3) as m^6^A-binding proteins to enhance the stability of thousands of cellular transcripts in an m^6^A-dependent manner [43]. It is possible that IGF2BPs bind to m^6^A-modified *PLIN3* mRNA and increase its stability in HIV-1 infected primary CD4^+^ T cells. However, despite this increase in mRNA levels, the level of PLIN3 protein was significantly decreased. The reason for this discrepancy is not clear. One possibility is that PLIN3 protein levels are regulated by mechanisms that are independent of post-transcriptional m^6^A modification. Alternatively, increased m^6^A modification of *PLIN3* mRNA may stabilize the transcript yet inhibit its translation. Further studies are needed to clarify these mechanisms.

Several previous studies explored the function of PLIN3 during HIV-1 infection. One study reported that HIV-1 Env binds to PLIN3 to target the trans-Golgi network in HeLa cells, which is essential for the efficient incorporation of HIV-1 Env into virions [44]. Subsequent studies built upon these observations and showed that PLIN3 interacts with HIV-1 Gag and Env [45], and is essential to produce infectious HIV-1 in both Jurkat T cells and primary macrophages [45, 46]. However, one group reevaluated the role of PLIN3 in HIV-1 infected Jurkat cells and found that PLIN3 was dispensable for Env virion incorporation [27]. However, whether PLIN3 plays a role in HIV-1 infection of primary CD4^+^ T cells has not been reported. In this study, we demonstrated that knockdown of PLIN3 in primary CD4^+^ T cells reduced HIV-1 virion release but increased virion infectivity. Since activated CD4^+^ T cells are the primary target of HIV-1 in vivo, these results are likely a more accurate reflection of how PLIN3 interacts with HIV-1 in the physiological environment. Considering the key role of plasma membrane rafts in HIV-1 assembly and release HIV-1 [42], it is plausible that PLIN3 affects the formation and function of plasma membrane rafts in primary CD4^+^ T cells, thereby regulating HIV-1 Gag and Env interaction during HIV-1 assembly and release. Future studies will focus on the role of PLIN3 in HIV-1 Env incorporation to better understand how PLIN3 regulates HIV-1 infection in primary CD4^+^ T cells.

In summary, here we report, at single-base resolution, m^6^A modification sites on viral and cellular RNA that occur in response to HIV-1 infection of primary CD4^+^ T cells. In addition, we have clarified previously conflicting results obtained in cell lines by establishing a role for PLIN3 in modulating HIV-1 infection in primary CD4^+^ T cells.

## Materials and Methods

### Ethics statement

The Institutional Review Board (IRB) at the University of Iowa has approved the in vitro experiments in this study involving human blood cells from de-identified healthy donors. The consent requirements for the de-identified blood samples were waived by IRB.

### Cell culture

Jurkat and primary CD4^+^ T cells were cultured in RPMI-1640 (ATCC) supplemented with 10% fetal bovine serum (FBS; R&D Systems) and antibiotics (100 U/mL penicillin and 100 µg/mL streptomycin, Gibco). HEK293T, Ghost/X4/R5, and TZM-bl cells were cultured in DMEM (Gibco) with 10% FBS and antibiotics [8]. All cells were cultured at 37°C with 5% CO_2_ and tested negative for mycoplasma contamination using a PCR-based universal mycoplasma detection kit (ATCC 30-1012K). Healthy deidentified donor blood was purchased from the DeGowin Blood Center at the University of Iowa. PBMCs were isolated from healthy donor blood as described (REF). CD4^+^ T cells were enriched using EasySep^™^ Human CD4^+^ T cell isolation kit (17952, STEMCELL Technologies) and activated using ImmunoCult^™^ Human CD3/CD28/CD2 T cell activator (10970, STEMCELL Technologies) for 72 hr.

### HIV-1 production and infection

Replication-competent HIV-1_NL4-3_ stocks were generated by transfection of HEK293T cells with pNL4-3 using jetPRIME (114-07, Polyplus Transfection) as described [8]. The supernatants were filtered (0.45 μm) and digested with DNase I (Turbo, Invitrogen) for 30 min at 37°C. The viral stock infectivities were calculated through serial dilution on Ghost/X4/R5 cell lines. For HIV-1 infection, Jurkat and primary activated CD4^+^ T cells were infected with HIV-1_NL4-3_ at an MOI of 1. Spinoculation was performed by centrifuging the cells with virus at 1200xg for 2 hr at 25°C. Cells were washed twice with Dulbecco’s Phosphate-Buffered Saline (DPBS) and resuspended with fresh culture medium. The reverse transcriptase inhibitor nevirapine (NVP, 10 μM, 4666, the AIDS Research and Reference Reagent Program, NIH) was used as a control. HIV-1 supernatant p24 levels were detected by p24 ELISA using anti-p24-coated plates (AIDS and Cancer Virus Program, National Cancer Institute, Frederick, MD) as described [47].

### Antibodies and immunoblotting

Antibodies used for immunoblotting were as follows: HIV-1 p24 (clone #24-2, the AIDS Research and Reference Reagent Program, NIH), GAPDH (AHP1628, Bio-Rad), METTL3 (15073, Proteintech), METTL14 (CL4252, Abcam), PLIN3 (10694-1-AP, Proteintech), HIV-Ig (3957, the AIDS Research and Reference Reagent Program, NIH). Cells were harvested and lysed in cell lysis buffer (9803, Cell Signaling Technology) with a protease and phosphatase inhibitor (A32959, Pierce, Thermo Scientific). Immunoblotting was performed as described [48]. GAPDH was used as a loading control for all immunoblots.

### RNA isolation and poly(A) enrichment

Total RNA was extracted using TRIzol (Invitrogen) and the RNA concentrations were determined by Nanodrop. mRNA was enriched using Dynabeads oligo(dT)25 (61005, Invitrogen) following the manufacturer’s instructions.

### m^6^A ELISA

m^6^A levels were quantified in 50 ng mRNA using a m^6^A RNA methylation ELISA protocol as described [13, 49].

### Co-IP assay

Cells were harvested and lysed in RIPA buffer (50 mM Tris-HCl (pH 7.5), 150 mM NaCl, 1% NP-40, 5 mM EDTA, and 10% glycerol) containing protease and phosphatase inhibitor. METTL3 complexes were precipitated with METTL3 antibody and Dynabeads™ protein G (1004D, Invitrogen). The same amounts of rabbit IgG were used as the negative control. The beads were washed three times with RIPA buffer and resuspended in LDS sample buffer (NP0007, Invitrogen). Input and IP samples were analyzed by immunoblot.

### m^6^A-SAC-Seq, data deposition, access, and bioinformatics analysis

-seq data have been m^6^A-SAC-seq was performed as previously described [15, 16]. Purified mRNA (150 ng) from each sample was used for m^6^A-SAC-seq. The m^6^A-SAC deposited in the Gene Expression Omnibus (GEO) with accession number GSE280563 (is scheduled to be released on May 01, 2025) https://www.ncbi.nlm.nih.gov/geo/query/acc.cgi?acc=GSE280563 The “pheatmap”, “ggplot2”, and “ggrepel” R package was used to identify differentiated m^6^A modifications on cellular genes, with thresholds set at fold change ≥ 2 with *p*<0.05. The “ggplot2” R package was employed to visualize GO and KEGG enrichment analyses using Metascape [50], with a threshold of *p*<0.05 indicating significant enrichment.

### meRIP

Total cellular RNA was isolated using TRIzol and its concentration were determined by Nanodrop. Total RNA was resuspended with IP buffer (50 mM Tris-HCl (pH7.5), 150 mM NaCl, 0.1% NP-40, and RNase Inhibitor). m^6^A antibody (202003, Synaptic Systems) or rabbit IgG (cat, vendor) were used for RNA-IP. The Monarch RNA Cleanup Kit (T2030S, New England Biolabs) was used to purify the enriched RNA. RT-PCR was conducted to detect target genes enrichment levels. Data analysis was performed using the ΔΔCt method.

### RT-PCR and quantitative PCR

Total RNA was extracted using Trizol or RNeasy Plus Kit (74134, Qiagen). cDNA was synthesized from the extracted RNA using iScript™ cDNA Synthesis Kit (1708891. Bio-Rad), and quantitative PCR (qPCR) was performed to quantify cDNA levels. Primers sequences are listed in Table S3.

### Plasmids

pLentiCRISPR v2 was from Feng Zhang (Addgene plasmid #52961). pLentiCRISPR v2 sgPLIN3 was constructed by ligating an oligonucleotide duplex (Integrated DNA Technologies) into the BsmBI-v2 site (R0739S, New England Biolabs). Oligonucleotide sequences used were listed in Table S3. Plasmids were confirmed by Sanger Sequencing.

### Generation of PLIN3 KO stable Jurkat cell lines and PLIN3 knockdown primary cells

Jurkat cells were transduced with lentiviruses in the presence of polybrene (10 µg/mL) by spinoculation at 1,200 x g for 2 hr at room temperature. Transduced cells were cultured in complete RPMI-1640 for 48 hr prior to selection with puromycin (1.5 µg/mL). After 7 days of selection, single-cell clones were obtained by limiting dilution. PLIN3 KO Jurkat cells were confirmed by immunoblotting and Sanger sequencing of genomic DNA. Primary CD4^+^ T cells were transduced with lentiviruses in the presence of polybrene (10 µg/mL) by spinoculation at 1,200 x g for 2 hr at room temperature. The transduced cells were then cultured in complete RPMI-1640 for 24 hr before undergoing a second round of transduction. After 48 hr transduction, the efficiency of PLIN3 knockdown was confirmed by immunoblotting.

### HIV-1 infectivity measurement in TZM-bl cell lines

TZM-bl cells (1×10^5^) were seeded in 24-well plates overnight and infected with 2 ng p24 HIV-1 stocks. After 48 hr, the luciferase activity was measured using ONE-Glo™ EX Luciferase Assay System (E8120, Promega). Luminescence was quantified using a microplate reader and normalized to total protein content.

### mRNA stability assay

Mock and HIV-1 infected primary CD4^+^ T cells were treated with actinomycin D (10 µg/mL) to inhibit transcription. Total RNA was extracted at 0, 1, 2, 4, and 6 hr post-treatment and qRT-PCR was performed to quantify the remaining *PLIN3* mRNA levels.

### Statistical analysis

Data were analyzed using *t*-test or analysis of variance (ANOVA) with Prism software and statistical significance was defined as *P* < 0.05.

## Acknowledgements

We thank the Wu lab members for helpful discussions and suggestions. We appreciate the reagents provided by the National Institutes of Health (NIH) AIDS Reagent Program. This work was supported by the NIH grants R61AI169659 to L. W. and RM1HG008935 to C.H. L. W. is also supported by NIH grants R01AI141495, R21AI170070, R21AI181742, and P30CA086862-24S1.

The content is solely the responsibility of the authors and does not necessarily represent the official views of the NIH. C.H. is an investigator of Howard Hughes Medical Institute.

## Author contributions

Conceptualization: Siyu Huang, Stacia Phillips, Li Wu

Resources: Li Wu, Chuan He

Methodology, Investigation, and Validation: Siyu Huang, Yutao Zhao, Bethany Wilms

Formal analysis: Siyu Huang, Stacia Phillips, Yutao Zhao, Li Wu

Writing – Original Draft: Siyu Huang, Stacia Phillips, Li Wu

Writing – Review & Editing: Siyu Huang, Stacia Phillips, Yutao Zhao, Chuan He, Li Wu

Visualization: Siyu Huang, Yutao Zhao, Stacia Phillips, Li Wu

Supervision, Project administration, and Funding acquisition: Chuan He, Li Wu

Library preparation and sequencing data analysis: Yutao Zhao, Chuan He

## Supporting information (Table S1-S3 in Excel files)

**Table S1. m^6^A-SAC-seq and RNA-seq data of primary CD4^+^ T cells.** Cells were infected with Mock or HIV-1 for 96 hr and poly(A)-enriched RNA were analyzed based on three individual healthy donors. m^6^A-SAC-seq and RNA-seq data are in two separate sheets in one Excel file. Data of genes listed in Table 1 are highlighted in red (m^6^A-SAC-seq).

**Table S2. m^6^A-SAC-seq identifies m^6^A modification sites in HIV-1 RNA in HIV-1 infected primary CD4^+^ T cells.** Cells were infected with Mock or HIV-1 for 96 hr and poly(A)-enriched RNA were analyzed based on three individual healthy donors. m^6^A motifs, m^6^A/A ratio of individual samples and their average values are included in the Excel file.

**Table S3. PCR primers sequences.**

